# Dopamine signaling modulates the stability and integration of intrinsic brain networks

**DOI:** 10.1101/252528

**Authors:** Golia Shafiei, Yashar Zeighami, Crystal A. Clark, Jennifer T. Coull, Atsuko Nagano-Saito, Marco Leyton, Alain Dagher, Bratislav Mišić

## Abstract

Dopaminergic projections are hypothesized to stabilize neural signaling and neural representations, but how they shape regional information processing and large-scale network interactions remains unclear. Here we investigated effects of lowered dopamine levels on within-region temporal signal variability (measured by sample entropy) and between-region functional connectivity (measured by pairwise temporal correlations) in the healthy brain at rest. The acute phenylalanine and tyrosine depletion (APTD) method was used to decrease dopamine synthesis in 51 healthy participants who underwent resting-state functional MRI (fMRI) scanning. Functional connectivity and regional signal variability were estimated for each participant. Multivariate partial least squares (PLS) analysis was used to statistically assess changes in signal variability following APTD as compared to the balanced control treatment. The analysis captured a pattern of increased regional signal variability following dopamine depletion. Changes in hemodynamic signal variability were concomitant with changes in functional connectivity, such that nodes with greatest increase in signal variability following dopamine depletion also experienced greatest decrease in functional connectivity. Our results suggest that dopamine may act to stabilize neural signaling, particularly in networks related to motor function and orienting attention towards behaviorally-relevant stimuli. Moreover, dopamine-dependent signal variability is critically associated with functional embedding of individual areas in large-scale networks.

## Introduction

The brain is a complex network of interacting neuronal populations that collectively support perception, cognition and action. Transient episodes of synchrony establish brief windows for communication among remote neuronal populations, manifesting as patterns of functional connectivity and large-scale resting state networks [25, 72, 91]. Thus, regional neural activity reflects computations that result from network interactions, but also drives those interactions [4, 26]. Greater connectivity may promote greater signal exchange, leading to variable dynamics [61, 78]; alternatively, densely interconnected regions may be more likely to synchronize, rendering their dynamics less variable and more stable [39]. How the balance between local dynamics and global functional interactions (connectivity) is modulated remains a fundamental question in systems neuroscience.

Dopamine is thought to stabilize neuronal signaling by modulating synaptic activity and signal gain [82]. Dopamine, acting in cortex or striatum, could regulate cortical representations by facilitating or suppressing neural signaling. These effects may also play a role in reinforcement learning, based on the theory of dopaminergic reward prediction error signaling [80]. In humans, transient decreases in dopamine synthesis (which we term *“dopamine depletion”*) have been shown to disrupt multiple aspects of perception, motor control and executive function [23, 66, 67, 73], consistent with a role in the regulation of sustained cortical activity [82]. Similar effects have also been demonstrated in various animal models including rodents and monkeys [81]. Furthermore, death of dopamine neurons in Parkinson’s disease (PD) leads to unstable and increasingly variable motor output [14, 52]. Thus, by stabilizing neuronal signaling, dopamine may influence the stability of regional activity and its potential for functional interactions at a network level.

Here we use resting-state functional magnetic resonance imaging (fMRI) to investigate the effects of dopamine depletion on within region signal variability and intrinsic brain networks in healthy brain at rest. We applied acute phenylalanine and tyrosine depletion (APTD) to transiently decrease dopamine levels in healthy participants [21, 50, 51, 57, 65, 69]. We hypothesized that dopamine depletion would destabilize regional hemodynamic activity, manifesting as increased signal variability. We further hypothesized that regions with increased signal variability may be less likely to interact with other regions, resulting in decreased functional connectivity defined by temporal statistical association of fMRI time series.

## Materials and methods

### Participants

Altogether, *n* = 51 healthy young individuals (righthanded, 23.6 *±* 5.9 years old, 32 male/19 female) participated in 3 separate dopamine precursor depletion studies (two published [23, 66] and one unpublished studies). The protocol, acquisition site, scanner and sequence were identical across the 3 studies. Participants with a history of drug abuse, neurological or psychiatric disorder were excluded. Informed consent was obtained from all participants.

### Dopamine depletion

The acute phenylalanine and tyrosine depletion (APTD) technique [51, 57, 69] was used to reduce dopamine synthesis in healthy participants, following the procedure described previously [23, 66]. In short, each participant was tested twice on two separate days, once following administration of a nutritionally balanced amino acid mixture (BAL) and once following acute phenylalanine/tyrosine depletion (APTD), in a randomized, double-blind manner, such that neither the participants nor the experiment conductors had any information regarding the label of the condition (BAL vs. APTD) being tested on each day. It should be noted that although APTD leads to depletion of dopamine precursors and only reduces the dopamine synthesis and availability, the term *“dopamine depletion”* is used throughout this manuscript to refer to *“dopamine precursor depletion”* and APTD. Although APTD might also theoretically decrease norepinephrine synthesis, several reports have shown that the release of norepinephrine is not affected under resting state conditions [48, 56].

### Data acquisition and preprocessing

T1-weighted, three-dimensional structural MRIs were acquired for anatomical localization (1-*mm*^3^ voxel size), as well as two resting-state echoplanar T2*-weighted images with blood oxygenation level-dependent (BOLD) contrast (3.5-mm isotropic voxels, TE 30 ms, TR 2 s, flip angle 90°) from all participants using a Siemens MAGNETOM Trio 3T MRI system at the Montréal Neurological Institute (MNI) in Montréal, Canada. Each participant was scanned for 5 minutes (150 volumes) with eyes open, on two separate days, once following administration of a nutritionally balanced amino acid mixture (BAL) and once following acute phenylalanine/tyrosine depletion (APTD). The resting state fMRI data was pre-processed through the following steps: slice timing correction, rigid body motion correction, removal of slow temporal drift using a high-pass filter with 0.01 Hz cut-off, physiological noise correction. Head motion parameters were estimated by spatially re-aligning individual time points with the median volume, which was then aligned with the anatomical T1 image of the individual. Further motion correction was done by scrubbing [71]: time points with excessive in-scanner motion (>0.5mm framewise displacement) were identified and removed from time series, along with the two volumes before and two volumes after. All preprocessing steps were performed using the Neuroimaging Analysis Kit (NIAK) [9, 10].

Anatomical MRI data was parcelled into 83 cortical and subcortical areas using the Desikan-Killiany atlas [28], and then further subdivided into 129, 234, 463 and 1015 approximately equally sized parcels following the procedure described by Cammoun and colleagues [20]. The Desikan-Killiany atlas is a commonly-used, anatomical (as opposed to functional), automated labeling system, where nodes are delineated according to anatomical landmarks. It has been shown that the Desikan-Killiany atlas is comparably reliable to manual parcellations of human cortex [28]. The atlas exists at 5 progressively coarser resolutions (the so-called ‘Lausanne’ parcellation [20]), allowing us to verify the experimental effects on various spatial scales. The parcellations were used to extract BOLD time series from functional MRI data. The time series of each parcel were estimated as the mean of all voxels in that parcel. All analyses were repeated at each resolution to ensure that none of the conclusions were idiosyncratic to a particular spatial scale.

### Sample entropy

Sample entropy (SE) analysis was used to estimate within-region signal variability [74]. The algorithm quantifies the conditional probability that any two sequences of time points with length of *m* + 1 will be similar to each other, given that the first *m* points of these sequences were similar (Fig. 1). SE is then measured as the natural logarithm of this quantity, such that large values are assigned to unpredictable signals, and small values to predictable signals. The algorithm is subject to two parameters: the pattern length (*m*), which determines the segment length used to find repeating patterns, and the similarity criterion (*r*), which is the tolerance for accepting matches in the time series. The sample entropy of a time series with length *N* is estimated as

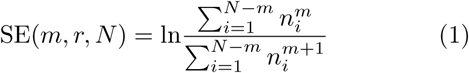

where 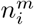 is the number of *m*-length segment of time series (e.g. segment *j* with length *m*) that are similar to the *m*-length segment *i* within to the similarity criterion, excluding self-matches (*i* ≠ *j*; i.e., the algorithm does not compare patterns with themselves) [22]. The sample entropy of a time series corresponds to ‘scale 1’ of the well-known multi-scale entropy analysis procedure [22].

**Figure 1.**
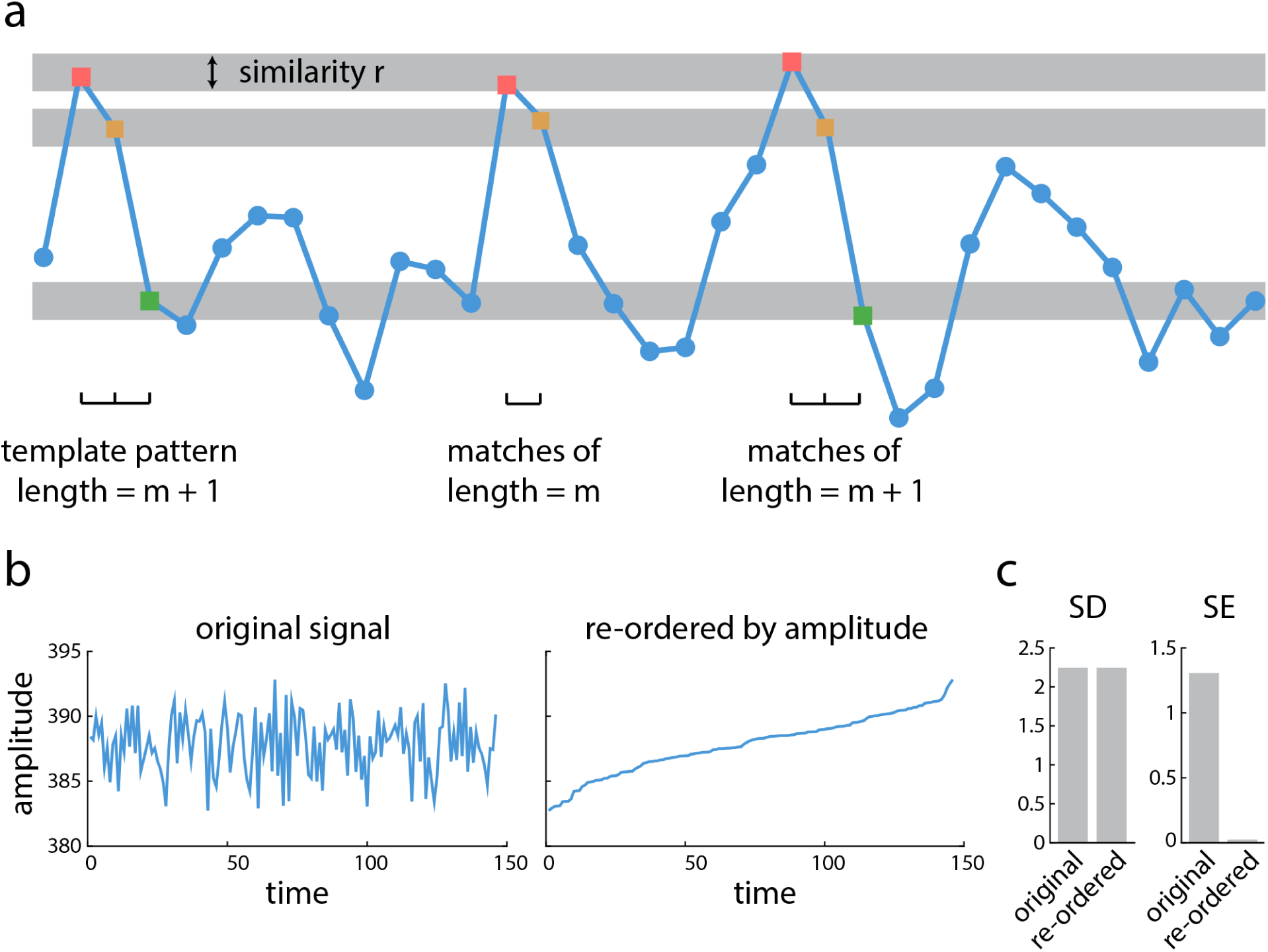
Sample entropy of a time series |. (a) An example of a BOLD signal is shown, where the x-axis is time and the y-axis is the amplitude. Signal variability is calculated using sample entropy analysis. Sample entropy (SE) measures the conditional probability that any two sequences of data points with length *m* + 1 will be similar to one another under the condition that they were similar for the first *m* points. The similarity criterion *r* represents the tolerance of algorithm to accept matches in the time series. (b) An example of a BOLD signal in its original form (left). The same signal, with the time points re-ordered by amplitude (right). (c) Standard deviation of the signal is the same for both the original and reordered signal; however, sample entropy of the re-ordered signal drastically decreases compared to sample entropy of the original signal.

Following the optimization proposed by Small & Tse [88], we set *m* = 2 as the pattern length. We set the similarity criterion to *r* = 0.5 times the standard deviation (SD) of the time series following the method proposed by Richman & Moorman [74]. Although these values of *m* and *r* have been used extensively in previous reports [8, 43, 53, 59], we sought to ensure that the reported results were robust across multiple choices of *m* and *r*. We therefore re-calculated SE using different values for *m* and *r* and re-ran the PLS analysis described below (see *Statistical assessment*). Fig. S2 shows the correlation between new bootstrap ratios (i.e., changes in signal variability) with bootstrap ratios that were originally estimated by setting *m* = 2 and *r* = 0.5 × *SD*. The correlations were generally greater than 0.7 across a range of similarity criteria *r*, and greater than 0.3 across a range of pattern lengths *m*, suggesting that the results were relatively insensitive to choice of parameters.

We operationalized signal variability using SE rather than other popular measures, such as standard deviation (SD). The primary reason for this choice is that SE is sensitive to temporal dependencies in the signal, while variance-based measures, such as SD, are not. This distinction is illustrated in Fig. 1b and Fig. 1c. Fig. 1b (left) shows a typical BOLD signal from the present study (a randomly selected condition, participant and node). Fig. 1b (right) shows the same signal, but with the time points re-ordered by amplitude. The sample entropy and standard deviation of the original and re-ordered signals were then measured (Fig. 1c). Sample entropy is sensitive to this change, because the re-ordered signal monotonically increases and is trivially predictable. Critically, standard deviation is blind to this change; although the temporal complexity of the signal has been profoundly altered by re-ordering, standard deviation measures only the dispersion of points and cannot detect any temporal change (Fig. 1c).

### Statistical assessment

We used partial least squares (PLS) analysis to investigate within-participant changes in regional signal variability following the BAL vs. APTD conditions. PLS analysis is a multivariate statistical technique that is used to analyze two “blocks” or sets of variables [54, 55]. In neuroimaging studies, one set may represent neural activity, while the other may represent behaviour or experimental design (e.g. condition and/or group assignments). PLS analysis seeks to relate these two data blocks by constructing linear combinations of the original variables such that the new latent variables have maximum covariance [46].

In the present report, one block (**X**) corresponded to regional signal variability in each participant estimated by sample entropy of BOLD time series following BAL vs. APTD conditions. The rows of matrix **X** correspond to observations (participants nested within conditions) and the columns correspond to variables (regional signal variability). For *p* participants, *c* conditions, and *v* variables, matrix **X** will have *p × c* rows and *v* columns. Within-condition means are computed for each column and centered to give the matrix **M**. Singular value decomposition (SVD) is applied to **M**

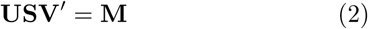

resulting in a set of orthonormal left singular vectors, **U** and right singular vectors, **V**, and a diagonal matrix of singular values, **S**. The number of latent variables is equal to the rank of the mean-centered matrix (here *c*), so **U** will have *c* columns and *v* rows, and **V** and **S** will both have *c* columns and *c* rows.

The decomposition results in a set of latent variables that are composed of columns of singular vectors, **U** and **V**, and a set of singular values from the diagonal matrix of **S**. In the present study, the *v* elements of column vectors of **U** are the weights of original brain activity variables (i.e., signal variability) that contribute to the latent variable and demonstrate a pattern of changes in signal variability following dopamine depletion. The *c* elements of column vectors of **V** are the weights of experimental design variables that contribute to the same latent variable and are interpreted as a contrast between experimental conditions. The latent variables are mutually orthogonal and express the shared information between the two data blocks with maximum covariance. This covariance is reflected in the singular values from the diagonal elements of matrix **S** that are associated with each given latent variable.

We assessed the statistical significance of each latent variable using permutation tests [30]. During each permutation, condition labels for each participant are randomized by reordering the rows of matrix **X**. The new permuted data matrices were then mean-centered and subjected to SVD as before. The procedure was repeated 10,000 times to generate a distribution of singular values under the null hypothesis that there is no relationship between neural activity and study design. A *p*-value was estimated for each latent variable as the proportion of permuted singular values greater than or equal to the original singular value.

We assessed the reliability of singular vector weights using bootstrap resampling. Here, the rows of data matrix **X** were randomly resampled with replacement while keeping the original condition assignments. Mean-centering and SVD were then applied to the resampled data matrices as before. The results were used to build a sampling distribution for each weight in the singular vectors **U** and **V**. A “bootstrap ratio” was then calculated for each original variable (i.e., for each node) as the ratio of the singular vector weight to its bootstrap-estimated standard error. Bootstrap ratios are designed to be large for variables that have a large weight (i.e., large contribution) as well as a small standard error (i.e., are stable). Bootstrap ratios are equivalent to z-scores if the bootstrap distribution is approximately unit normal [31]. In this case, 95% and 99% confidence intervals correspond to bootstrap ratios of ±1.96 and ±2.58, respectively.

PLS was chosen as the primary analytic method (instead of univariate statistical techniques) because we sought to identify patterns of nodes whose signal variability collectively changes due to dopamine depletion. However, the results of PLS analysis were nearly identical with the results obtained by a more conventional univariate paired t-test (correlation between t-values and bootstrap ratios; *r* ≈ 1).

### Community detection

Functional networks were partitioned into communities or intrinsic networks using the assignment derived in [62], which we describe below. As we show in the *Results* section, the main conclusions also hold for the partitions reported by Yeo and colleagues [91] and Power and colleagues [72].

A Louvain-like greedy algorithm was used to identify a community assignment that maximized the quality function, *Q* [68, 77]

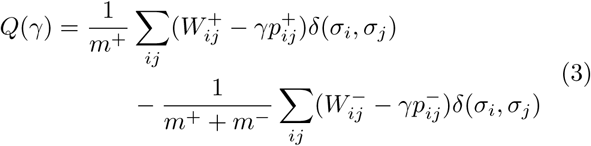

*where* 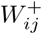 and 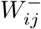 are the functional connectivity (correlation) matrices that contain only positive and only negative coefficients of correlation, respectively. 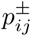 is the expected density of only positive or only negative connectivity matrices according to the configuration null model and is given as 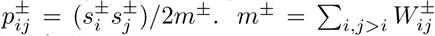 is the total weight of all positive or negative connections of 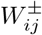 (note that the summation is taken over *i, j > i* to ensure that each connection is only counted once). The total weights of positive or negative connections of *i* and *j* are given by 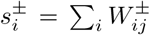 and 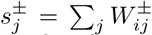, respectively. The resolution parameter *γ* scales the importance of the null model and effectively controls the size of the detected communities: larger communities are more likely to be detected when *γ* < 1 and smaller communities (with fewer nodes in each community) are more likely to be detected when *γ* > 1. Furthermore, *σ*_*i*_ defines the community assignment of node *i*. The Kronecker function *δ*(*σ*_*i*_*, σ*_*j*_) is equal to 1 if *σ*_*i*_ = *σ*_*j*_ and is equal to zero otherwise (*σ*_*i*_ ≠ *σ*_*j*_), ensuring that only within-community connections contribute to *Q*.

Multiple resolutions *γ* were assessed, from 0.5 to 10 in steps of 0.1. The Louvain algorithm was repeated 250 times for each *γ* value [15]. The resolution *γ* = 1.5 was chosen based on the similarity measures (Rand index) of pairs of partitions for each *γ* value, such that the similarity measures of a more stable set of partitions for a given *γ* value would have a larger mean and smaller standard deviation compared to similarity measures at other *γ* values (i.e., larger z-score of similarity measure) [90]. Finally, a consensus partition was found from the 250 partitions at *γ* = 1.5 following the method described in [6]. Eight communities or networks were detected, including visual (VIS), temporal (TEM), default mode (DMN), dorsal attention (DA), ventral attention (VA), somatomotor (SM) and salience (SAL) [62]. The subcortical areas (SUB) were added to the list as a separate network based on the anatomical Desikan-Killiany parcellation.

### Cohesion and integration

Connectivity between and within modules was assessed as the participation coefficient and within-module degree z-score [41], using Brain Connectivity Toolbox (BCT) [76]. The participation coefficient quantifies how evenly distributed a node’s connections are to all modules. Values close to 1 indicate that a node is connected to many communities, while values close to 0 indicate that a node is connected to few communities. The participation coefficient of node *i*, *P*_*i*_, is given by:

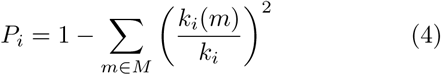

where *m* is a module from a set of modules *M*, *k*_*i*_ is the weighted degree (i.e., connection strength) of node *i*, and *k*_*i*_ (*m*) is the number of connections between node *i* and all other nodes in module *m* [41, 76]. To find participation coefficients of resting state networks, we first found the average participation coefficient of each node across all subjects and then compared the participation coefficients of the nodes that belong to the same module following the BAL vs. APTD conditions.

The within-module degree is estimated as the weighted degree (i.e., strength) of the connections that node *i* makes to other nodes within the same module. The measure is then z-scored, expressing a node’s weighted degree in terms of standard deviations above or below the mean degree of the nodes in the given module (*Z*_*i*_). A positive within-module degree z-score indicates that a node is highly connected to other nodes within the same module, while a negative within-module z-score indicates a node participates in less than average connections within its own module. We estimated the within-module degree z-score of each node for each subject and then calculated the average *Z*_*i*_ over all subjects. Finally, we compared the within-module degree z-scores of the nodes of a given module following the BAL and APTD conditions.

## Results

Task-free, eyes-open resting-state fMRI was recorded in *n* = 51 healthy young participants on two separate days, once following administration of a nutritionally balanced amino acid mixture (BAL) and once following acute phenylalanine/tyrosine depletion (APTD). Anatomical MRI data were parcelled into five progressively finer resolutions, comprising 83, 129, 234, 463 and 1015 nodes [20]), which were used for extraction of blood-oxygen-level dependent (BOLD) time series. We investigated how dopamine depletion affects (a) local, region-level hemodynamic activity, (b) global, between-region temporal statistical association of BOLD time series (termed as *functional connectivity*) and (c) the relationship between the two.

### Dopamine depletion increases signal variability

We estimated within region signal variability using sample entropy (SE), a measure of the unpredictability of a time series [74]. Briefly, the SE algorithm quantifies the conditional probability that any two sequences of *m* + 1 time points will be similar to each other given that the first *m* points were similar (Fig. 1). We then used multivariate partial least squares (PLS) analysis to statistically assess within-participant changes in signal variability at each brain region following administration of the BAL vs. APTD mixtures [55]. PLS results in a set of latent variables (LV), that are weighted combinations of experimental design (i.e., a contrast) and signal variability patterns that optimally covary with each other.

The analysis revealed a single statistically significant latent variable (permuted *p* = 0.014 for the finest parcellation resolution with 1015 nodes), showing broadly increased signal variability following dopamine depletion (Fig. 2). Bootstrap resampling was used to estimate the reliability with which individual nodes contribute to the overall multivariate pattern. Specifically, the weight or loading associated with each node was divided by its bootstrap-estimated standard error, yielding a measure (“bootstrap ratio”) that is high for nodes with large weights (i.e., large contributions) and small standard errors (i.e., are stable) [54]. Note that bootstrap ratios may be interpreted as z-scores if the sampling distribution is approximately unit normal [31]. Positive bootstrap ratios indicate an increase in signal variability, while negative bootstrap ratios indicate decreased variability. Fig. 2c depicts a brain projection of this statistical pattern, showing that the greatest increase in signal variability was observed in somatomotor cortex. This effect (increased regional hemodynamic variability following depletion) and the spatial pattern were consistent across all five spatial resolutions (Fig. S1).

**Figure 2.**
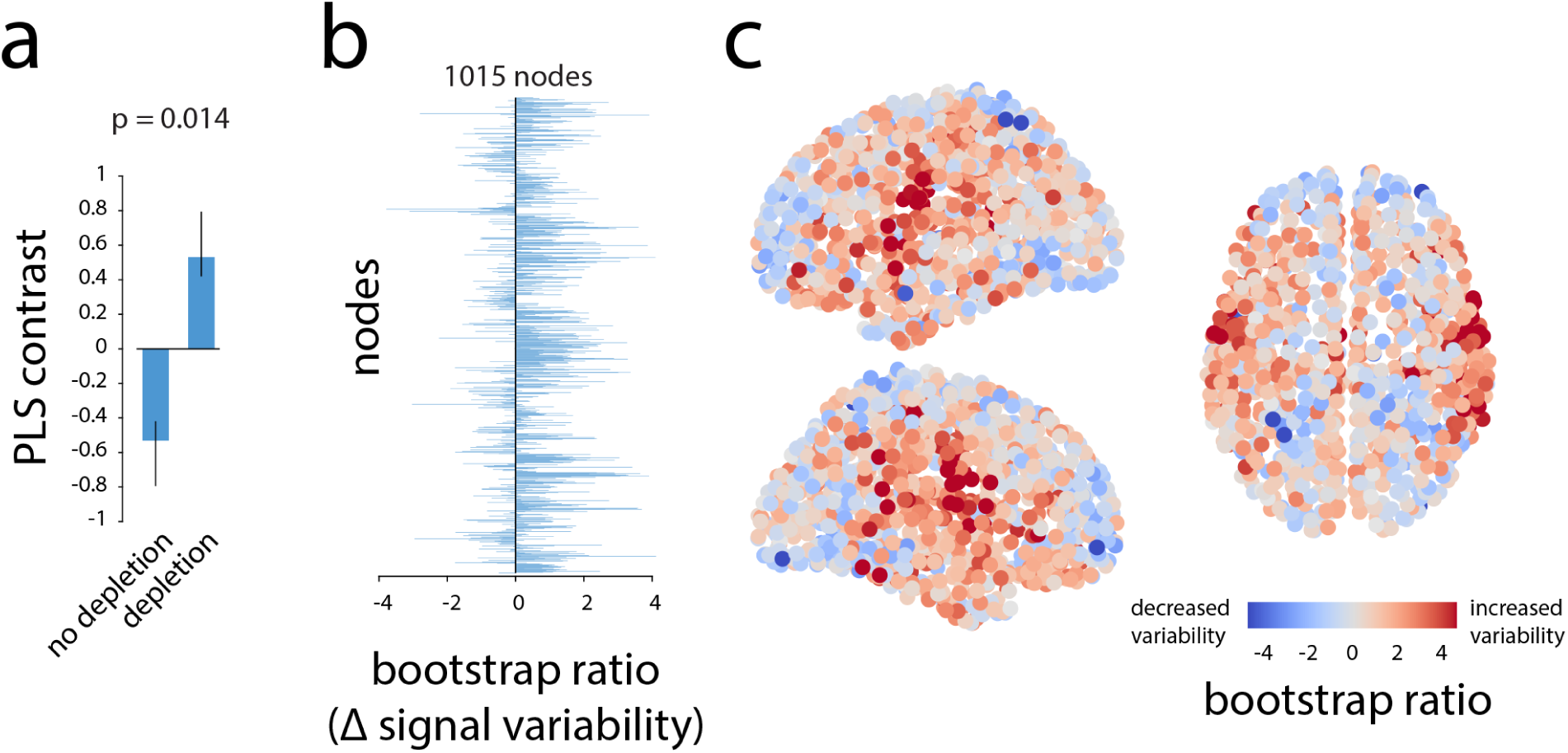
Dopamine depletion increases signal variability |. (a) PLS analysis identified a significant contrast between patterns of signal variability in depletion (APTD) vs. non-depletion (BAL) conditions (permuted *p* = 0.014). (b) The change in signal variability of each node is given by a bootstrap ratio for that node: such that a positive bootstrap ratio shows increase in signal variability of the node following dopamine depletion, while a negative bootstrap ratio shows the opposite. Bootstrap ratios are depicted at the finest resolution (1015 nodes), showing that dopamine depletion increases signal variability at most nodes. (c) Bootstrap ratios are shown in 3D space sagittally and axially. Corresponding results are shown for all resolutions in Fig. S1

### Increased signal variability in somatomotor and salience networks

We next investigated the effect of APTD on resting state networks [72, 91]. Fig. 3a depicts the nodes displaying the greatest increase in signal variability following dopamine depletion in descending order and colour-coded by resting state network membership [62]. The most affected nodes appear to belong primarily to the somatomotor (yellow) and salience (green) networks suggesting that the signal variability may selectively affect certain large-scale networks.

**Figure 3.**
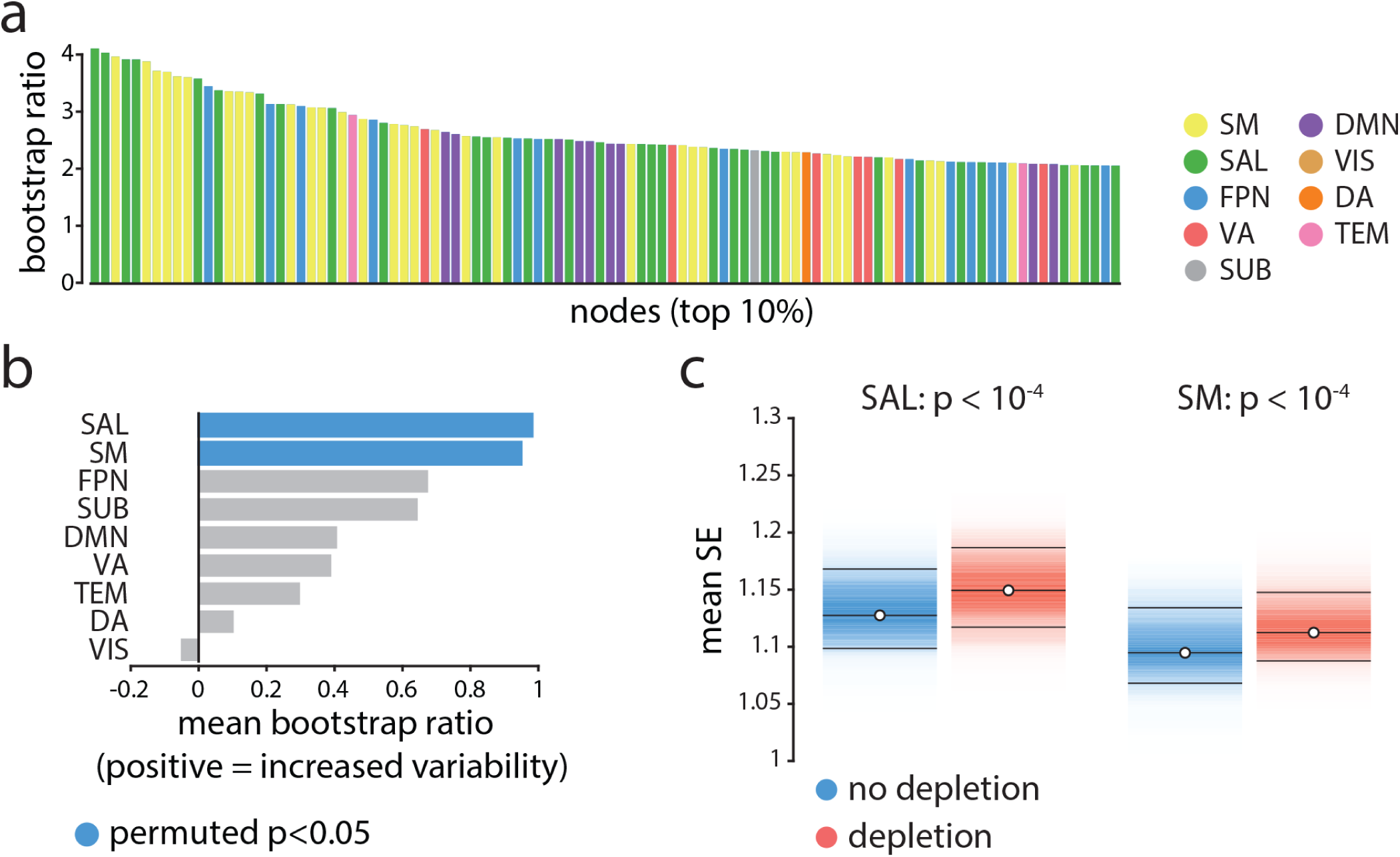
Nodeand network-level effects of dopamine depletion |. (a) The top 10% of the nodes (i.e., top 100 nodes) that had the largest increase in signal variability (largest bootstrap ratios) following dopamine depletion. Each bar shows the magnitude of bootstrap ratio of a node and is colored based on the community assignment of that node [62]. Somatomotor (yellow) and salience (green) networks appear over-represented compared to other networks. (b) The mean change in signal variability is calculated for each network and assessed by permutation tests (10,000 repetitions). Signal variability increases most in the salience and somatomotor networks following dopamine depletion, and these are the only two networks where this effect is statistically significant. (c) Changes in mean signal variability are depicted for somatomor and salience networks (significance obtained by permutation tests; FDR corrected [11]). SM = somatomotor, SAL = salience, FPN = fronto-parietal, VA = ventral attention, SUB = subcortical areas, DMN = default mode, VIS = visual, DA = dorsal attention, TEM = temporal.

To directly investigate the network-selectivity of dopamine depletion, we first estimated the mean change in signal variability across all nodes in a given network, using PLS-derived bootstrap ratios for the finest resolution (1015 nodes). To determine which network-level changes were statistically significant and not due to differences in network size, spatial contiguity or lateralization, we used a label permuting procedure. Network labels were randomly permuted within each hemisphere (preserving the number of nodes assigned to each network) and network means were recomputed 10,000 times, generating a distribution under the null hypothesis that network assignment does not influence the overall change in signal variability. A *p*-value was estimated for each network as the proportion of cases when the mean for the permuted network assignment exceeded the mean for the original empirical network assignment. Fig. 3b-c shows that changes in signal variability were observed for all intrinsic networks, but that increased variability was greatest and statistically significant for the somatomotor and salience networks (*p* < 10^−4^, FDR corrected [11]).

To ensure that these results are independent of how intrinsic networks are defined, we repeated the procedure using partitions reported by Yeo and colleagues [91] and by Power and colleagues [72] (Fig. S3). The results were consistent across the three partitions, indicating significant increased signal variability in somatomotor and ventral attention networks among Yeo networks (note that the ‘ventral attention network’ overlaps with the ‘salience network’ shown in Fig. 3), and in somatosensory and auditory networks among Power networks. No significant decrease in signal variability was observed in any intrinsic networks, regardless of which network assignments were used.

### Increased signal variability correlates with decreased functional connectivity

Given that changes in signal variability were highly network dependent, we next investigated whether increased signal variability is related to patterns of functional connectivity. Functional connectivity was estimated as a zero-lag Pearson correlation coefficient between regional time series for each participant in each condition. To relate patterns of signal variability with functional connectivity, we estimated a group-average functional connectivity matrix by calculating the mean connectivity of each pair of brain regions across all participants. We then estimated the mean connectivity (i.e., strength of the functional correlations) for each brain region.

We observed a weak relationship between increased signal variability and decreased functional connectivity, such that nodes with the greatest increase in signal variability following dopamine depletion also experienced the greatest decrease in functional connectivity (*r* = –0.23, *R*^2^ = 0.053, *p* = 1.04 *×* 10^−13^; Fig. 4a). Although statistically significant, effect was small, suggesting that the relationship was not sustained over the bulk of data points (nodes), but that it may have been driven by a subset of nodes instead.

**Figure 4.**
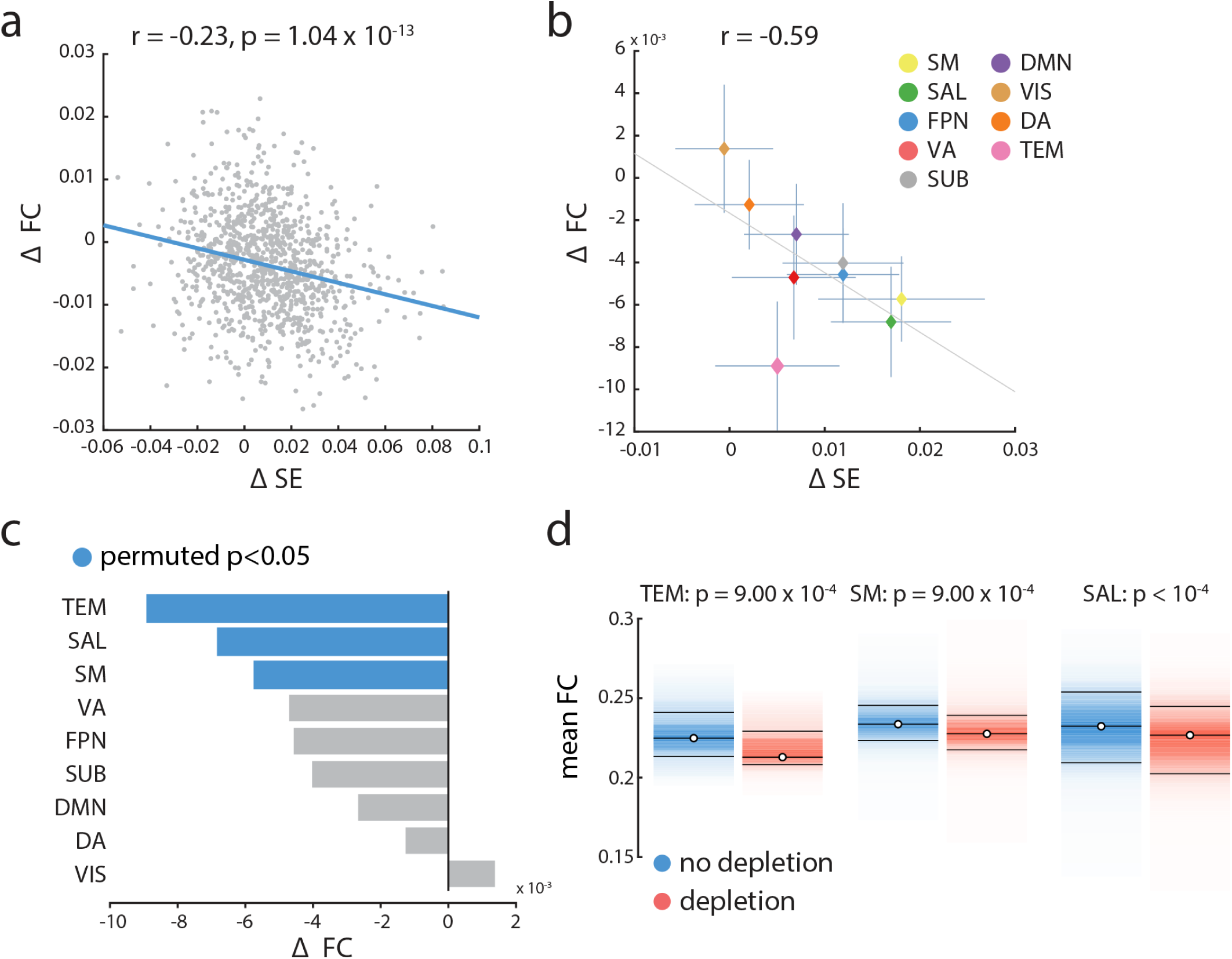
Relating signal variability and functional connectivity |. (a) Mean changes in functional connectivity following dopamine depletion were estimated across all nodes and correlated with changes in within-region signal variability. Changes in functional connectivity are related to changes in signal variability, such that the larger the increase in signal variability, the larger the decrease in functional connectivity. (b) Mean changes in functional connectivity for intrinsic networks are correlated with mean changes in local signal variability in those networks. There is a clear anti-correlation between the two, consistent with the result in part (a). (c) The mean changes in functional connectivity was calculated for each network and assessed by permutation tests (10,000 repetitions). Mean connectivity significantly decreases in temporal, salience and somatomotor networks. Somatomotor and salience networks also experience significant increase in local variability (Fig. 3). (d) Mean functional connectivity in depletion (APTD) vs. non-depletion (BAL) conditions, shown for nodes belonging to the temporal (TEM), somatomotor (SM) and salience (SAL) networks. Functional connectivity decreases in all instances (permutation test; FDR corrected).

To investigate this possibility, we assessed dopamine-dependent changes in functional connectivity in each of the intrinsic networks separately and correlated the network specific changes in functional correlation with the network specific changes in signal variability. We observed an anti-correlation between the two measures such that the networks with greatest increase in hemodynamic signal variability also experience the great decrease in functional correlations (*r* = –0.59; Fig. 4b). Changes in functional connectivity were statistically assessed using the same label permuting procedure outlined above (randomly permuting the network label of all nodes and re-computing network means, with 10,000 repetitions). Mean functional connectivity significantly decreased in three intrinsic networks: temporal, salience and somato-motor networks connectivity (*p* = 9.0 *×* 10^−4^, *p <* 10^−4^ and *p* = 9.0 *×* 10^−4^ respectively; FDR corrected; Fig. 4c-d). Critically, the salience and somatomotor networks also experienced the greatest increase in signal variability after APTD (Fig. 3), suggesting that changes in signal variability and functional connectivity may be related. Overall, these results suggest that the effects of dopamine depletion are stronger in specific large-scale systems, and that changes in local dynamics are related to global functional interactions.

### Selective disconnection of intrinsic networks

Dopamine-related increases in signal variability appear to be concomitant with decreased functional connectivity localized to specific intrinsic networks. However, it is unclear whether decreased connectivity in the somatomotor and salience networks is driven by weakened within-network or between-network connections, or both. To address this question, we calculated the participation coefficient and within-module degree z-score of every node [41]. The participation coefficient quantifies the diversity of a node’s connectivity profile. A participation coefficient with a value close to 1 indicates that a node’s connections are evenly distributed across communities, while a value close to 0 indicates that most of the node’s connections are within its own community. The within-module degree z-score of a node is a normalized measure of the strength of connections a node makes within its own community.

Fig. 5 shows that the participation coefficient significantly decreases in somatomotor and salience networks following dopamine depletion (*p* < 10^−4^ and *p* = 2.4 *×* 10^−3^ respectively, assessed by label permuting (see above); FDR corrected), while the within-module degree z-score does not (*p* > 0.5). In other words, dopamine depletion selectively reduced functional interactions between these networks and the rest of the brain (participation coefficient; Fig. 5a), but did not affect within-network cohesion (within-module degree z-score; Fig. 5b). Overall, these results suggest that dopamine depletion effectively segregates these intrinsic networks from the rest of the brain, but does not affect their internal cohesion. Note that the two networks with significantly decreased participation coefficient are also the ones with greatest increases in signal variability. We also investigated average changes in participation coefficient and within-module degree z-score in other intrinsic networks, where we did not observe any significant changes in either of the two measures.

**Figure 5.**
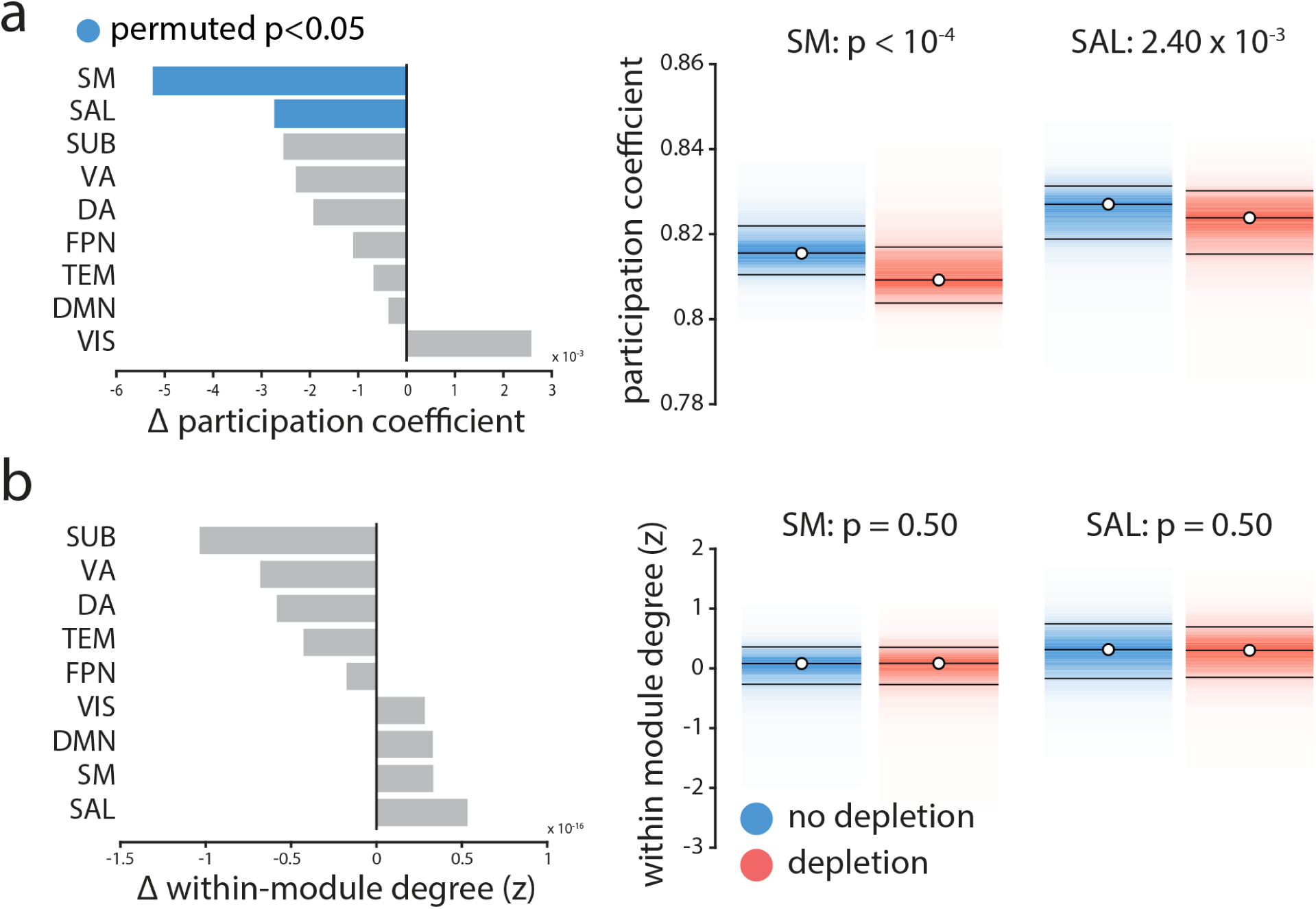
The effects of dopamine depletion on cohesion and integration of specific intrinsic networks |. (a) Mean participation coefficient, indexing the diversity of inter-network connectivity, significantly decreases in somatomotor (SM) and salience (SAL) networks after dopamine depletion (using 10,000 permutation tests; FDR corrected). (b) Within-module degree z-score, indexing within-network connectivity, remains unaffected.

### No systematic effect of study

The data used in this study were consolidated from three different experiments (two published [23, 66] and one unpublished studies), so it is possible that the observed effects were idiosyncratic to one or two of the constituent datasets and do not necessarily generalize across all three studies. To investigate this possibility, we used a multi-way ANOVA to assess differences between studies: subject-specific scores were calculated for the signal variability pattern and the three studies were treated as separate groups. The analysis did not reveal any significant difference among the three studies (*F* (2, 45) = 1.7, *p* = 0.19), nor any condition by study interaction (*F* (2, 45) = 0.68, *p* = 0.51). There was a significant condition difference, with greater scores in the depletion versus non-depletion condition (*F* (1, 45) = 131.06, *p* ≈ 0), but this is expected given that the scores were derived by PLS to maximize this condition difference.

### Comparing sample entropy and standard deviation

A popular alternative measure of signal variability is the simple standard deviation (SD). Although we opted to use SE over SD because the latter is not sensitive to temporal dependencies in the signal (see Fig. 1), for completeness we directly compared the effects of depletion using the two measures. A priori, we expect the two measures to be anti-correlated, because sample entropy estimation explicitly incorporates the standard deviation of a given signal to define the similarity criterion *r* (the tolerance of the algorithm to accept matches in the time series). In other words, the similarity criterion for the sample entropy algorithm will be greater for a signal with a greater standard deviation. Consequently, the sample entropy algorithm is more likely to identify matches in signals with a larger SD, resulting in a lower sample entropy value. To demonstrate this claim, we correlated the standard deviation and sample entropy of regional time series for both APTD (depletion) and BAL (no depletion) conditions, as well as the changes in each measure following dopamine depletion. The results are shown in Fig. S4, confirming the anti-correlation between sample entropy and standard deviation of a given signal at a brain region in the (a) no depletion condition, (b) depletion condition. Panel (c) further shows that depletion-driven changes in signal variability are also anti-correlated with changes in standard deviation.

## Discussion

We investigated the effect of dopamine depletion on the balance between local node properties and global network architecture. We report two key results: (1) dopamine depletion selectively destabilizes neural signaling” measured at the hemodynamic level, in salience and somatomotor networks, and (2) increased local variability in these intrinsic networks is accompanied by their disconnection from the global functional architecture. Altogether, these results point to a stabilizing influence of dopamine on neural signaling and highlight the link between local, node-level properties and global network architecture.

### Linking local and global dynamics

The present results highlight the relationship between local hemodynamic signal variability and functional embedding. Increased variability in salience and somatomotor networks was concomitant with decreased functional connectivity with the rest of the brain. It is possible that low dopamine states disrupt local neuronal signaling, making it less likely for remote populations to synchronize. Alternatively, dopamine depletion may disrupt inter-regional synchrony through a separate mechanism, resulting in greater local variability. The correlative nature of the results cannot be used to disambiguate these two possibilities and further causal experiments are necessary. In any case, the present report demonstrates that functional interactions span multiple topological scales, such that local and global dynamical properties cannot be fully appreciated in isolation [16, 18].

Interestingly, dopamine depletion reduced between-module connectivity but did not affect within-module connectivity (Fig. 5). The effect was highly specific: reduced between-module connectivity was significantly observed only in networks that also experienced increased regional signal variability. In other words, dopamine depletion affected how nodes within these networks communicated with the rest of the brain, but did not affect their internal cohesion. A recent study demonstrated a similar effect at the level of resting state networks: networks with greater temporal variability displayed greater within-network cohesion and lower between-network integration [49]. Altogether, the present results highlight a simple principle: the tendency for nodes to form functional networks depends on their ability to synchronize with one another. Thus, functional interactions between regions must be studied together with the temporal properties of their local signals.

Recent theories emphasize dynamic over static brain function. At the network level, reconfiguration of functional interactions is increasingly recognized as an informative attribute of healthy brain function and dysfunction [19]. Functional reconfiguration has been observed across multiple temporal scales, both at rest [13, 92] and with respect to a variety of cognitive functions [86], including learning [7, 63], attention [87] and working memory [45], and even conscious awareness [5, 38]. In parallel, the dynamic range of local signal fluctuations has emerged as a node-level marker of brain function [36, 75]. Traditionally disregarded as “noise”, changes in signal variability have been reported across the lifespan [33, 42, 53], in multiple perceptual, cognitive and affective tasks [34, 60, 79] and in a variety of psychiatric and neurological diseases [12, 58, 59].

While most methods for estimating variability focus on node-level time series, several recent studies have conceptualized variability with respect to functional network embedding [59, 84]. For instance, local variability can be defined as the tendency for a node to switch network allegiance or to interact with multiple networks [17, 93]. This dynamic network switching is conditioned by an underlying anatomy [83, 93], but is also likely to be influenced by a variety of neurotransmitters. A prominent hypothesis is that dopamine modulates signal-to-noise ratio [64, 79]. We turn to the specific role of dopamine next.

### Dopamine and signal dynamics

Our results suggest that dopamine may act to stabilize neural signaling at the hemodynamic level, particularly in networks associated with motor control (somatomotor network) and orienting attention towards behaviorally-relevant stimuli (salience network). Dopamine depletion was simultaneously associated with increased signal variability and decreased extrinsic connectivity, indicating that dopaminergic signaling influences both local information processing and network-wide interactions. Importantly, the effects of depletion were not confined to a single locus but distributed over two large-scale networks, suggesting that even temporary decreases in dopamine availability can disrupt local neuronal signaling and have far-reaching effects on synchrony among multiple systems.

There are two possible mechanisms by which dopamine depletion could cause the observed changes in cortical signal variability. The first possibility is that depletion modulates synaptic activity and signal gain directly via cortical receptors [82]. Mechanistic studies *in vitro* have demonstrated that dopamine influences intrinsic ionic currents and synaptic conductance [29, 47]. These modulatory effects may facilitate or suppress neural signaling, helping to stabilize neural representations. In addition, dose-response effects of dopamine release may be both tonic and phasic [40], with the two modes thought to be mediated by distinct signaling pathways and receptors, and manifesting in distinct behavioural outcomes [24]. For instance, striatal medium spiny neurons of the direct pathway express *D*_1_ receptors and are thought to promote movement and the selection of rewarding actions. Neurons in the indirect pathway mainly express *D*_2_ receptors and are thought to inhibit cortical patterns that encode maladaptive or non-rewarding actions [89]. Although our results are consistent with the broad notion that dopamine stabilizes neural representations to facilitate reward learning and movement, further experiments are necessary to determine whether the observed effects can be attributed to tonic or phasic modulation, and to *D*_1_ or *D*_1_ receptor transmission.

The second possibility is that the effects of dopamine depletion may originate in the striatum, an area with dense dopaminergic afferents as well as projections to both the somatomotor and salience networks [2, 3, 94]. Prominent projections from dorsal striatum terminate in the somatomotor system (forming the motor loop), while projections ventral striatum terminate in the salience system. A dopaminergically-depleted striatum may therefore disrupt ongoing cortico-striatal signaling, resulting in downstream cortical effects, such as increased variability. Importantly, the two accounts are not mutually exclusive, and it is possible that the observed effects depend on both mechanisms.

Dopamine depletion can thus have local and global consequences, influencing a range of sensory-motor and higher cognitive functions. Age-related decline in dopaminergic transmission is hypothesized to lead to greater signal variability, influencing the distinctiveness of neural representations and, ultimately, performance [64, 79]. The stabilizing role of dopamine can also be observed in diseases associated with dopaminergic dysfunction, such as Parkinson’s disease (PD), attention deficit hyperactivity disorder (ADHD) and schizophrenia. In PD for instance, cell death in substantia nigra leads to reduced dopaminergic transmission, with extensive motor symptoms. Intriguingly, dopamine depletion in PD is associated with reduced cortico-striatal functional connectivity patterns and reduced gait automaticity [37]. Similarly, in ADHD, reduced dopamine signaling is associated with deficits in goal-directed behaviour and reward learning [27].

Finally, the present results draw attention to an overlooked assumption of graph-based models of brain structure and function: that all nodes are identical, except for their connectivity patterns. In other words, graph representations often ignore important inter-regional differences that could influence neural activity and synchrony, including morphology, cytoarchitectonics, gene expression and receptor densities. How dopaminergic modulation interacts with modulation by other neurotransmitters is an exciting open question [85].

### Measuring signal variability

Finally, we note that several recent reports have also investigated the role of dopaminergic signaling in the context of local signal dynamics, but drew an altogether different conclusion: that dopamine up-regulation increases signal variance. Specifically, Alavash and colleagues (2018) [1] reported that L-dopa administration increased BOLD standard deviation during an auditory working memory task (a syllable pitch discrimination task). Similarly, Garrett and colleagues (2015) [35] reported that d-amphetamine administration also increased signal standard deviation during a working memory task (a visual letter n-back task). Although we used a different method to manipulate dopamine (APTD vs. L-dopa and damphetamine) and to record hemodynamic activity (resting state vs. task), we believe that the primary difference between these studies and our own is how signal variability was operationalized. Namely, both Alavash et al. (2018) [1] and Garrett et al. (2015) [35] defined signal variability in terms of standard deviations. The results shown in Fig. 1 and Fig. S4 demonstrate that sample entropy and variance based measures (e.g., standard deviation) capture different aspects of signal variability. Most importantly, because of the way that sample entropy is used to detect repeating patterns in a signal, we find that in practice, the two measures are often anticorrelated, which explains the seemingly different results. Altogether, these studies demonstrate a need to further refine the concept of signal variability and for greater plurality of methods [32]. While some measures are sensitive to signal dispersion (e.g. standard deviation), others are sensitive to temporal regularity (e.g. sample entropy).

### Methodological considerations

Our results may depend on a number of methodological choices and potential limitations, which we consider in detail here. Methodological choices include the type of parcellation and resolution, intrinsic network definition, and parameter settings for SE. The reported effects are consistent across five resolutions (from 83 to 1015 nodes; Fig. S1), three network partitions (detected using clustering, Infomap and Louvain methods; Fig. S3) and a range of parameter settings (Fig. S2). Although we took steps to mitigate concerns about these choices, the present results are based on a finite sampling of a multifactorial methodological space.

More generally, we studied the effects of dopamine depletion in the context of task-free, resting state fMRI, which presents three significant challenges for interpretation. First, dopaminergic transmission is inherently related to specific cognitive functions, which may be accessible without overt task demands. We find evidence that dopamine depeltion affects information transfer in two intrinsic networks with specific functional properties, but more research is necessary to investigate how dopamine affects the function of these networks in the presence of task demands. Second, dopaminergic transmission within specific subcortical and cortical circuits occurs at time scales that may be inaccessible with BOLD imaging. The present results can be used to draw conclusions about slow, modulatory effects of dopamine, but more electrophysiological evidence is necessary to relate these effects to faster phasic dopaminergic responses. Third, the present data were collected during an eyes-open resting state scan, which may potentially entail different neurocognitive demands than eyes-closed, including recruitment of visuomotor and attention networks [44, 70].

### Summary

Our results support a link between local node dynamics and network architecture. Pharmacological perturbation may selectively target and disconnect specific networks without altering their internal cohesion. These results demonstrate that the stabilizing effect of dopamine on synaptic signaling extends to the level of large-scale brain networks.

## Acknowledgments

This research was undertaken thanks in part to funding from the Canada First Research Excellence Fund, awarded to McGill University for the Healthy Brains for Healthy Lives initiative. BM acknowledges support from the Natural Sciences and Engineering Research Council of Canada (NSERC Discovery Grant RGPIN #017-04265) and from the Fonds de recherche du Québec Santé (Chercheur Boursier). GS acknowledges support from the Healthy Brains for Healthy Lives (HBHL) initiative at McGill University. The authors gratefully acknowledge code from Dr. Richard F. Betzel (University of Pennsylvania, PA, USA) and Dr. Andrea Avena-Koenigsberger (Indiana University, IN, USA) to create the boxplot and scatter plot figures, respectively.

## Author Contributions

Conceptualization, G.S., B.M. and A.D.; Methodology and Resources, C.A.C, J.T.C., A.N.-S., M.L., A.D.; Data Curation, Y.Z.; Formal Analysis and Investigation, G.S., Y.Z.; Writing-Original Draft, G.S., B.M. and A.D.; Writing-Review & Editing, G.S., Y.Z., M.L., B.M. and A.D.; Funding Acquisition, M.L., A.D. and B.M.

---

**Figure S1.**
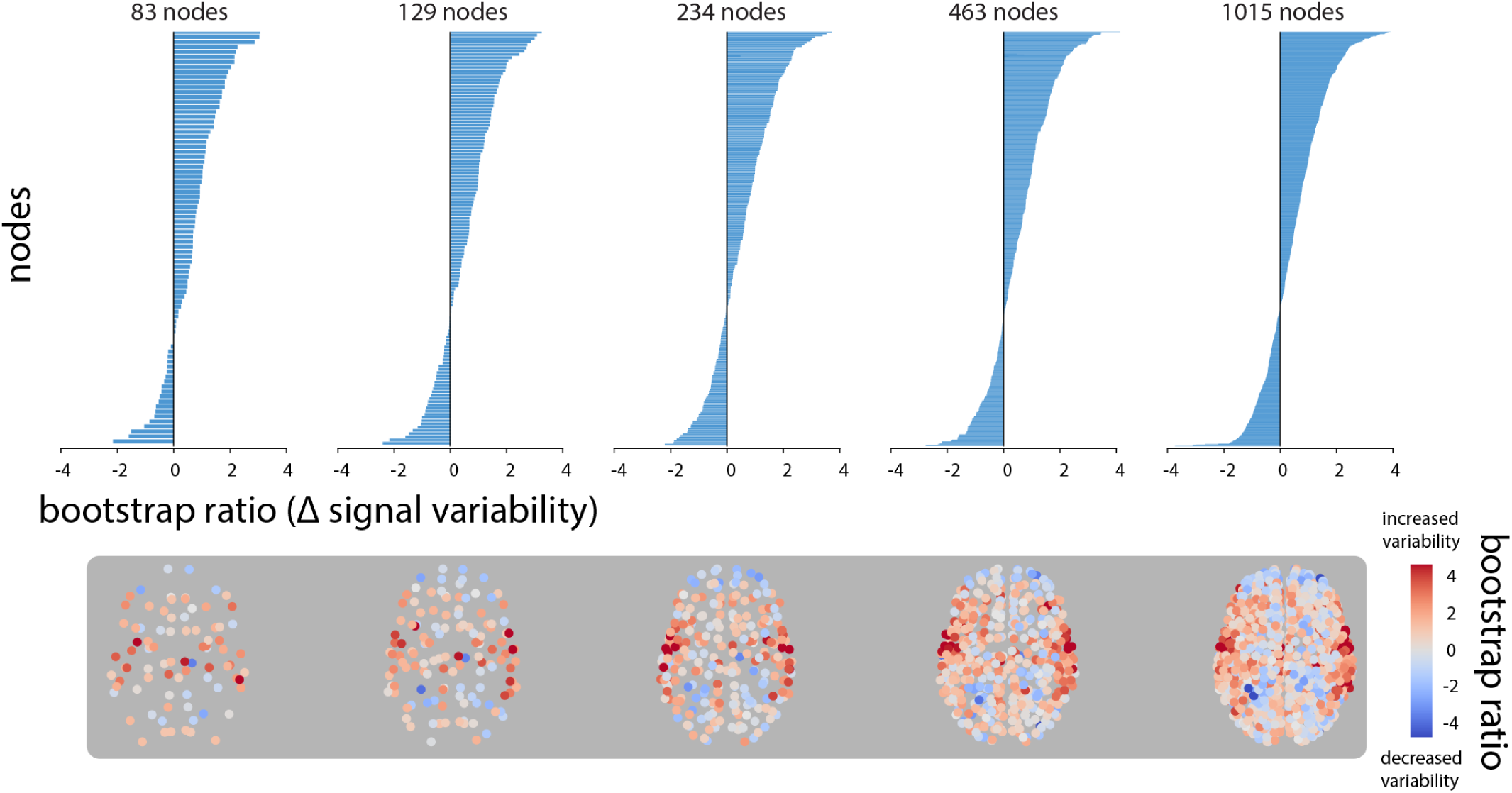
Replicating results using alternative parcellation resolutions |. In all parcellation resolutions, PLS analysis identifies a contrast between the patterns of signal variability in depletion (APTD) vs. non-depletion (BAL) conditions, (permuted *p*-values for the 5 resolutions from 83 nodes to 1015 nodes are as following: *p* = 0.047, *p* = 0.023, *p* = 0.006, *p* = 0.013, and *p* = 0.014). Changes in signal variability of each node is given by a bootstrap ratio for that node, such that a positive bootstrap ratio shows increased variability of the node following dopamine depletion, while a negative bootstrap ratio shows the opposite. The bootstrap ratios are ordered and depicted at all resolutions by bar graphs (first row) and on each node of the brain network (second row).

**Figure S2.**
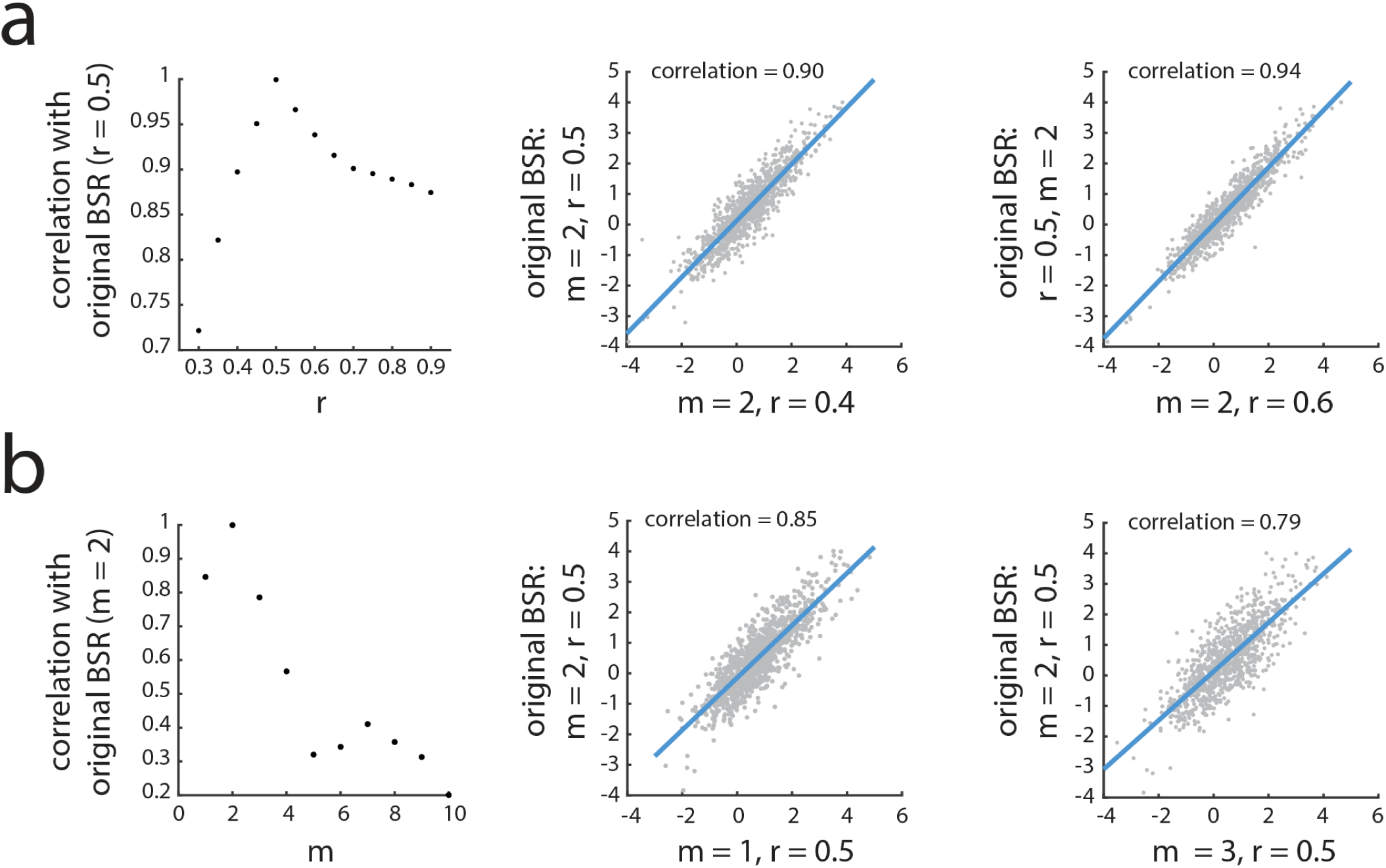
Choosing parameters for sample entropy analysis |. The whole analysis was repeated several times to ensure that results are not affected by the choice of parameters used to estimate signal variability. For this purpose, signal variability was calculated for various *m* and *r* values. Results of each *m* and *r* value were subjected to PLS analysis and new bootstrap ratios were estimated each time. (a) New bootstrap ratios estimated for varying similarity criterion, *r*, were correlated with original bootstrap ratios that were estimated with *r* = 0.5 × *SD*. The correlation coefficient at each *r* is shown in this figure. Note that pattern length was kept constant as *m* = 2. The correlation of bootstrap ratios for *r* = 0.4 × *SD* and *r* = 0.6 × *SD* with the original bootstrap ratios (*r* = 0.5 × *SD*) are shown as examples. (b) Keeping similarity criterion unchanged at *r* = 0.5 × *SD*, the correlation coefficients of new bootstrap ratios for varying pattern length, *m*, with original bootstrap ratio with *m* = 2 were estimated. Correlation of bootstrap ratios for *m* = 1 and *m* = 3 with original bootstrap ratios (*m* = 2) are depicted to show examples of varying *m*.

**Figure S3.**
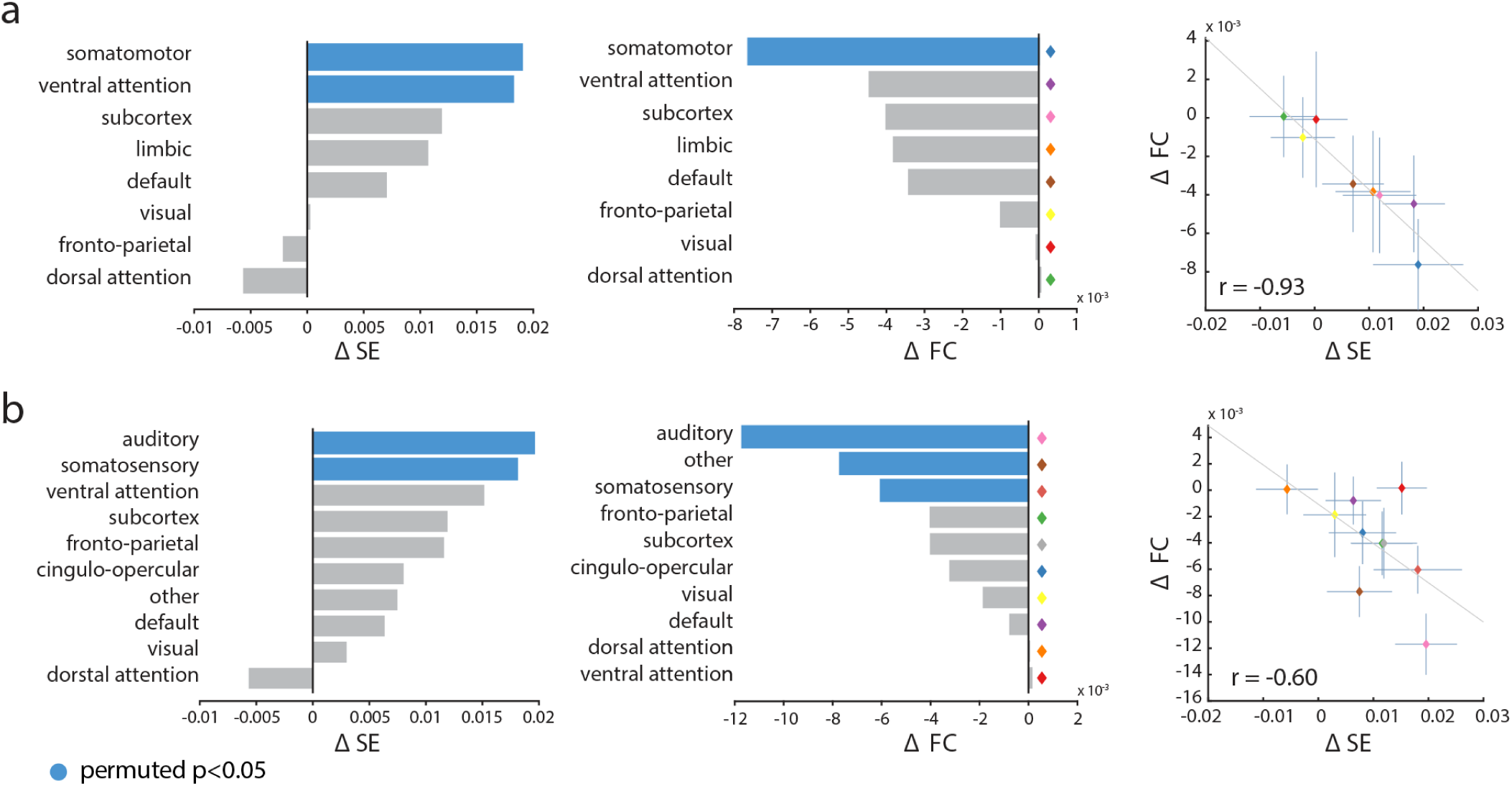
Replicating results using alternative network partitions |. Mean local signal variability and mean functional connectivity across nodes were estimated at each resting state brain network using two of other well-known community assignments defined by (a) Yeo and colleagues [91], and (b) Power and colleagues [72]. The significance of the mean signal variability and functional connectivity in each network was determined by 10,000 permutation tests. Mean network-wise changes in functional connectivity and local variability were correlated (scatter plots; *r* = –0.93 in (a) and *r* = –0.60 in (b)). The results are consistent with Fig. 3 and Fig. 4, confirming that our analysis is independent from the methods used to identify community assignments.

**Figure S4.**
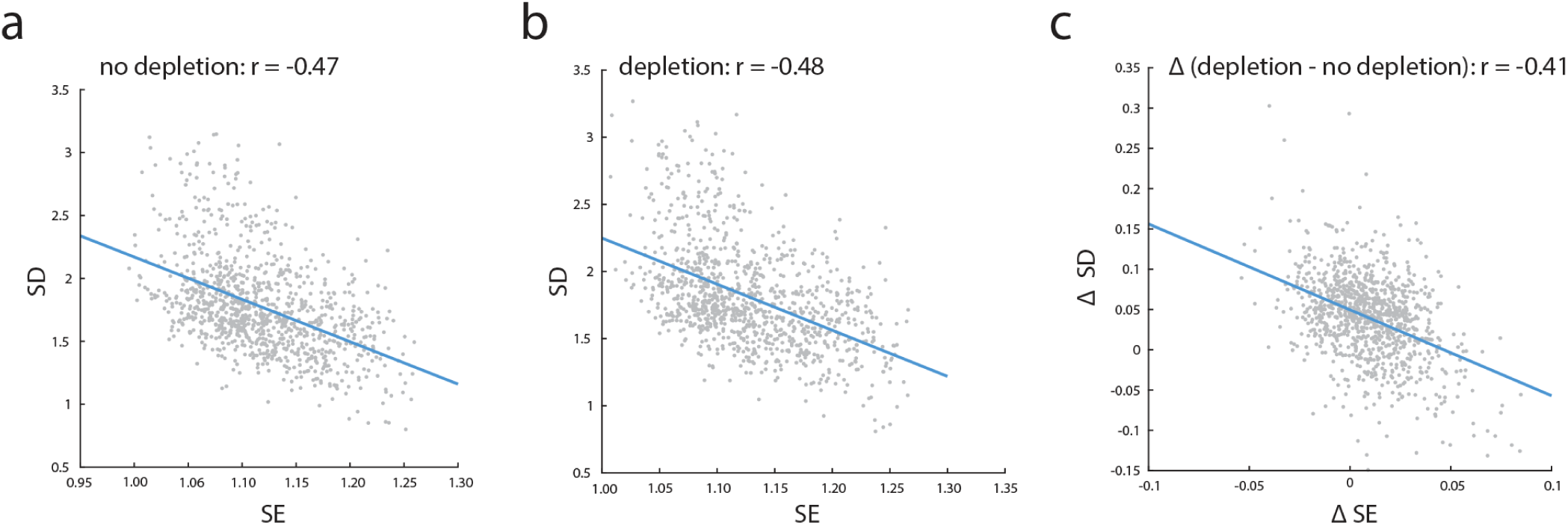
Sample entropy versus standard deviation |. Sample entropy (SE) and standard deviation (SD) of the BOLD signal at each brain region were calculated and correlated for (a) BAL (no dopamine depletion) and (b) APTD (dopamine depletion) conditions. SE and SD are anti-correlated in both conditions, such that larger SE of the BOLD signal in a brain region corresponds to smaller SD. (c) Changes in sample entropy (?SE) and standard deviation (?SD), following dopamine depletion, are also anti-correlated.

**Figure S5.**
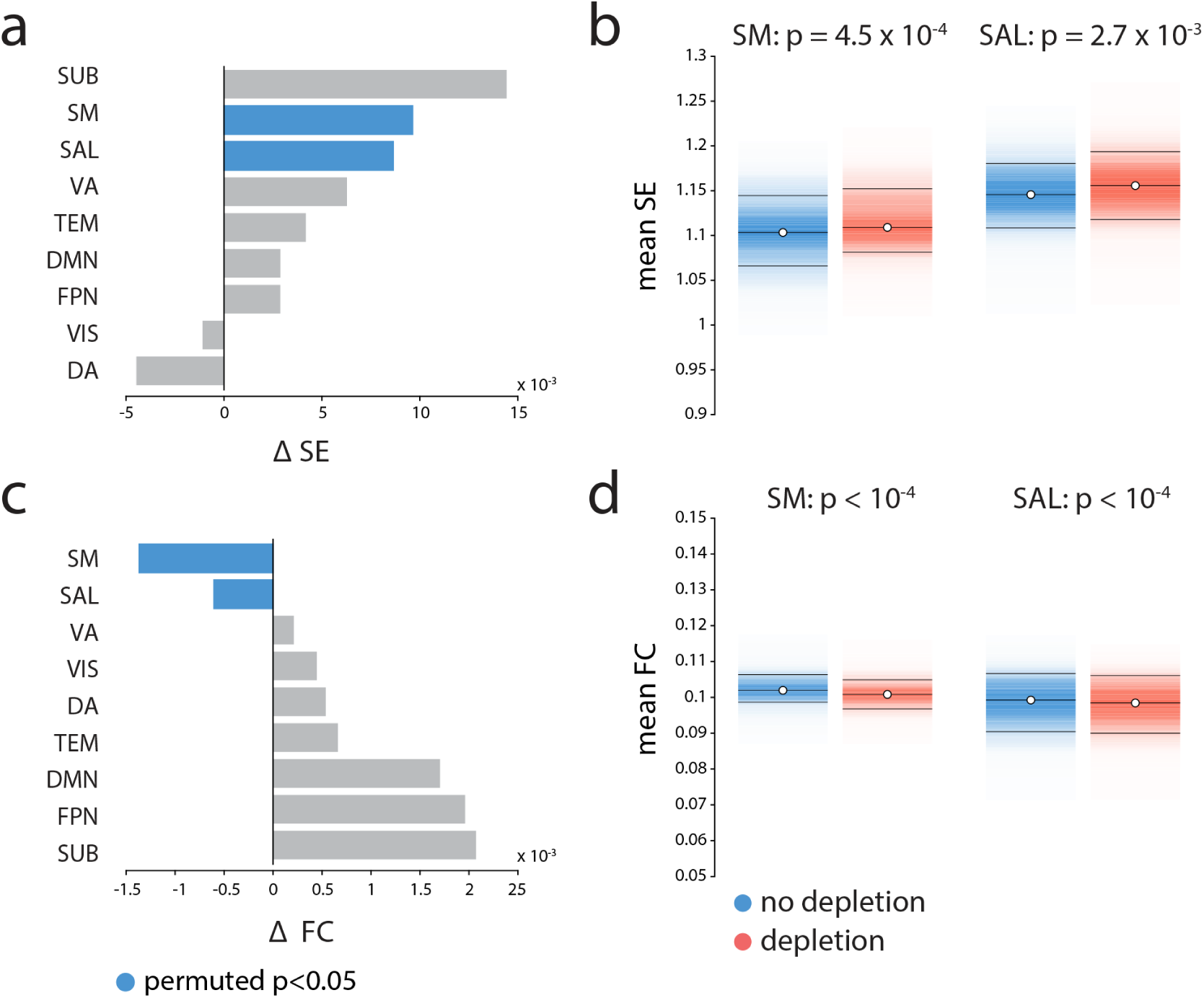
Replicating results after performing global signal regression |. The analysis was repeated following global signal regression. The results are consistent with the ones shown before: (a) Mean signal variability increases significantly in the somatomotor and salience networks following dopamine depletion. (b) Changes in mean signal variability are depicted for somatomor and salience networks (significance obtained by permutation tests; FDR corrected). (c) Mean functional connectivity significantly decreases in somatomotor and salience networks following dopamine depletion. (d) Changes in mean functional connectivity are depicted for somatomor and salience networks (significance obtained by permutation tests; FDR corrected). SM = somatomotor, SAL = salience, FPN = fronto-parietal, VA = ventral attention, SUB = subcortical areas, DMN = default mode, VIS = visual, DA = dorsal attention, TEM = temporal.

